# A Src family kinase maintains latent sensitization in rats, a model of inflammatory and neuropathic pain

**DOI:** 10.1101/2020.02.18.954859

**Authors:** Wenling Chen, Juan Carlos Marvizón

## Abstract

Latent sensitization is a long-term model of chronic pain in which hyperalgesia is continuously suppressed by opioid receptors. This is demonstrated by the induction of mechanical allodynia by opioid antagonists. Different intracellular signals may mediate the initiation, maintenance and expression of latent sensitization. Our criterion for the involvement of a signal in the maintenance of latent sensitization is that it inhibitors should permanently eliminate the allodynia produced by an opioid antagonist. We hypothesized that Src family kinases (SFKs) maintain latent sensitization and tested this hypothesis in rats with latent sensitization induced by complete Freund’s adjuvant (CFA) or by spared nerve injury. After measures of mechanical allodynia returned to baseline, the SFK inhibitor PP2 or vehicle were injected intrathecally. The opioid antagonist naltrexone injected intrathecally 15 min later produced allodynia in control rats but not in rats injected with PP2. PP2 or vehicle were injected daily for two more days and naltrexone was injected five days later. Again, naltrexone induced allodynia in the control rats but not in the rats injected with PP2. Results were similar when latent sensitization was induced either with CFA or spared nerve injury. We concluded that an SFK, likely Fyn, maintains latent sensitization induced by inflammation or nerve injury.

**Perspective:** This article presents evidence that a Src family kinase, likely Fyn, maintains latent sensitization induced by inflammation or nerve injury. If latent sensitization is a valid model of chronic pain, inhibiting its maintenance with Src family kinase inhibitors may cure chronic pain.

## Introduction

Chronic pain is a dysregulation of pain processing mechanisms that causes them to stay sensitized even after the healing of the initial injury. Evidence for this idea is provided by the latent sensitization model of chronic pain, which shows that certain injuries put the organism in a long-term state in which pain sensitivity is present but is continuously suppressed by opioid and α_2A_ adrenergic receptors (Taylor and Corder, 2014; Marvizon et al., 2015). The simultaneous presence of pain sensitization and its suppression is evidenced by the ability of antagonists of µ, δ and κ opioid receptors and α_2A_ adrenergic receptors to induce episodes of mechanical allodynia and thermal hyperalgesia (Campillo et al., 2011; Corder et al., 2013; Walwyn et al., 2016). Moreover, mice with genetic deletion of µ-opioid receptors, either global (Walwyn et al., 2016) or restricted to nociceptive afferents (Severino et al., 2018), are unable to fully recover from hyperalgesia and do not show induction of allodynia by an opioid antagonist.

There may be different forms of latent sensitization depending on its triggering injury: inflammation (Corder et al., 2013; Walwyn et al., 2016), nerve injury (Solway et al., 2011), plantar incision (Rivat et al., 2009; Campillo et al., 2011) or opioids (Rivat et al., 2007; Rivat et al., 2009). It needs to be determined whether they have different mechanisms.

We propose that different intracellular signals mediate the initiation, expression and maintenance of latent sensitization. Initiation signals would be those elicited by the triggering injury that leads to the establishment of latent sensitization. Maintenance signals would be those that ensure that the sensitized state endures over time. Expression signals would be those required for the occurrence of hyperalgesia when they are recruited by the maintenance signals. Drugs that inhibit the expression latent sensitization would eliminate the allodynia induced by opioid antagonists only while they are present, whereas drugs that inhibit its maintenance should eliminate antagonist-induced allodynia permanently. It is important to find drugs that inhibit the maintenance of latent sensitization because they may cure chronic pain by reversing pain mechanisms back to the normal state.

NMDA receptors (Celerier et al., 2000; Corder et al., 2013), adenylyl cyclase (Corder et al., 2013) and protein kinase A (Fu et al., 2019) are required for latent sensitization, but it is unclear whether they mediate its expression or its maintenance. NMDA receptors in the dorsal horn are potentiated by phosphorylation by the Src family kinase (SFK) Fyn (Guo et al., 2002; Abe et al., 2005; Chen et al., 2010) downstream of the activation of trkB receptors by BDNF (Mizuno et al., 2003; Xu et al., 2006; Geng et al., 2010; Chen et al., 2014; Li et al., 2017). Therefore, we hypothesized that an SFK mediates the maintenance of latent sensitization. We tested this hypothesis by determining whether PP2, an inhibitor of Fyn and other SFKs (Mizuno et al., 2003; Li et al., 2017; Berrout and Isokawa, 2018) reverses the allodynia produced by the opioid antagonist naltrexone (NTX) in latent sensitization induced by inflammation with complete Freund’s adjuvant (CFA) or by spared nerve injury (SNI).

## Methods

### Animals

Animals were male adult (2-4 months old) Sprague-Dawley rats (Envigo, Indianapolis, IN). A total of 30 rats were used in the study: 15 rats were injected in the hindpaw with saline or CFA for the first experiment and 15 rats underwent sham surgery or SNI for the second experiment. All animal procedures were approved by the Institutional Animal Care and Use Committee of the Veteran Affairs Greater Los Angeles Healthcare System, and conform to NIH guidelines. Efforts were made to minimize the number of animals used and their suffering.

### Chemicals

Naltrexone (NTX) and PP2 were purchased from Tocris Bioscience (Minneapolis, MN). CFA and other reagents were obtained from Sigma-Aldrich (St. Louis, MO). PP2 was dissolved at 10 mM in DMSO and then diluted to 30 µM in saline (final DMSO was 0.3%). NTX was dissolved in saline.

### CFA injections

Undiluted CFA was injected subcutaneously in a volume of 50 µl into one hindpaw using a 50 µl Hamilton syringe and a 26 gauge needle. The needle was inserted at an oblique angle from the heel in the middle of the paw, near the base of the third toe. It was held in place for 15 s and then gently withdrawn.

### Spared nerve injury (SNI) by common peroneal and sural transection

To reduce the initial period of allodynia, we used a variation of the SNI model in which the common peroneal and the sural branches of the sciatic nerve are cut (CpxSx) instead of the common peroneal and the tibial branches. After CpxSx, responses to von Frey filaments return to baseline in about 28 days (Solway et al., 2011). Rats were anesthetized with isoflurane. At the dorsal upper thigh level, the skin was cut and the muscle separated to expose the trifurcation of the sciatic nerve. The common peroneal and sural nerves were ligated and then cut on both sides of the suture to remove 2-3 mm of each nerve, leaving the tibial nerve intact. The muscle fascia were sutured in a simple continuous pattern. The cut in the skin was closed using a subcuticular intradermal pattern. Rats were given daily injections of the analgesic carprofen for 3 days. The presence of motor weakness or signs of paresis was established as a criterion for immediate euthanasia, but this did not occur in any of the rats.

### Intrathecal (i.th.) cannulation and injections

Rats were implanted with chronic intrathecal catheters under isoflurane (2–4%) anesthesia as described (Storkson et al., 1996; Walwyn et al., 2016). After cutting the skin and muscle, a 20G needle was inserted between the L5 and L6 vertebrae to puncture the dura mater. The needle was removed and the catheter (20 mm of PE-5 tube heat-fused to 150 mm of PE-10 tube) was inserted into the subdural space and pushed rostrally to terminate over L5-L6. The PE-10 end of the catheter was tunneled under the skin and externalized over the head. The skin was sutured and the catheter was flushed with 10 µl saline and sealed. Rats were housed separately and used 7 days after surgery. Intrathecal injection volume was 10 µl of injectate plus 10 μl saline flush. Solutions are preloaded into a PE-10 tube and delivered within 1 min. The position of the catheter was examined postmortem. Termination of the catheter inside the spinal cord or its occlusion were exclusion criteria, but this was not observed in any of the rats. Three rats lost the catheter after day 28 and were excluded from further measures.

### I.th. doses of NTX and PP2

NTX and PP2 were injected i.th. to restrict their effects to the spinal cord (Taylor and Corder, 2014). The dose of NTX (2.6 nmol i.th.) was the same used in previous studies (Corder et al., 2013; Taylor and Corder, 2014; Walwyn et al., 2016). The dose of PP2 was calculated based on its IC_50_ of 5 nM for Fyn (Lawrence and Niu, 1998). To obtain the i.th. dose from the IC_50_, it was multiplied by 30 to get a saturating concentration and then divided by 500 nM/nmol. This number was obtained as the ratio of the EC_50_s for NMDA to induce substance P release in spinal slices [258 nM (Chen et al., 2010)] and intrathecally *in vivo* [0.49 nmol (Chen et al., 2014)]. The resulting dose of PP2 was 0.3 nmol.

### Measures of mechanical allodynia

Mechanical allodynia was measured using von Frey filaments (Touch-Test) by the two-out-of-three method as described (Kingery et al., 2000; Michot et al., 2012; Jarahi et al., 2014; Walwyn et al., 2016; Chen et al., 2018). Rats were habituated for periods of 30 min for 3 days to acrylic enclosures on an elevated metal grid (IITC Life Science Inc., CA). A series of von Frey filaments were applied in ascending order to the plantar surface of the hindpaw for a maximum of 3 s. A withdrawal response was counted only if the hindpaw was completely removed from the platform. Each filament was applied three times, and the minimal value that caused at least two responses was recorded as the paw withdrawal threshold (PWT). The 15 g filament was taken as the cut-off threshold.

### Timeline

The timeline of the experiments is illustrated in Fig. 1. After habituating the rats to the von Frey enclosures for 3 days, i.th. catheters were surgically implanted on day −7. Baseline von Frey measures were taken on days −4 and 0. After the measure on day 0, rats received CFA or SNI in one hindpaw. The mechanical allodynia produced by CFA or SNI was followed with von Frey measures on days 3 and 21. On day 28, a baseline von Frey measure was taken and the rats received i.th. injections of vehicle or PP2, and i.th. NTX 15 min later. For the following 2 h, von Frey measures were taken at 15, 30, 60, 120 min after NTX. The i.th. injections of vehicle or PP2 were repeated on days 29 and 30. On day 35, a baseline von Frey measure was taken, NTX was injected i.th., and von Frey measures were taken for 2h.

**Figure 1.**
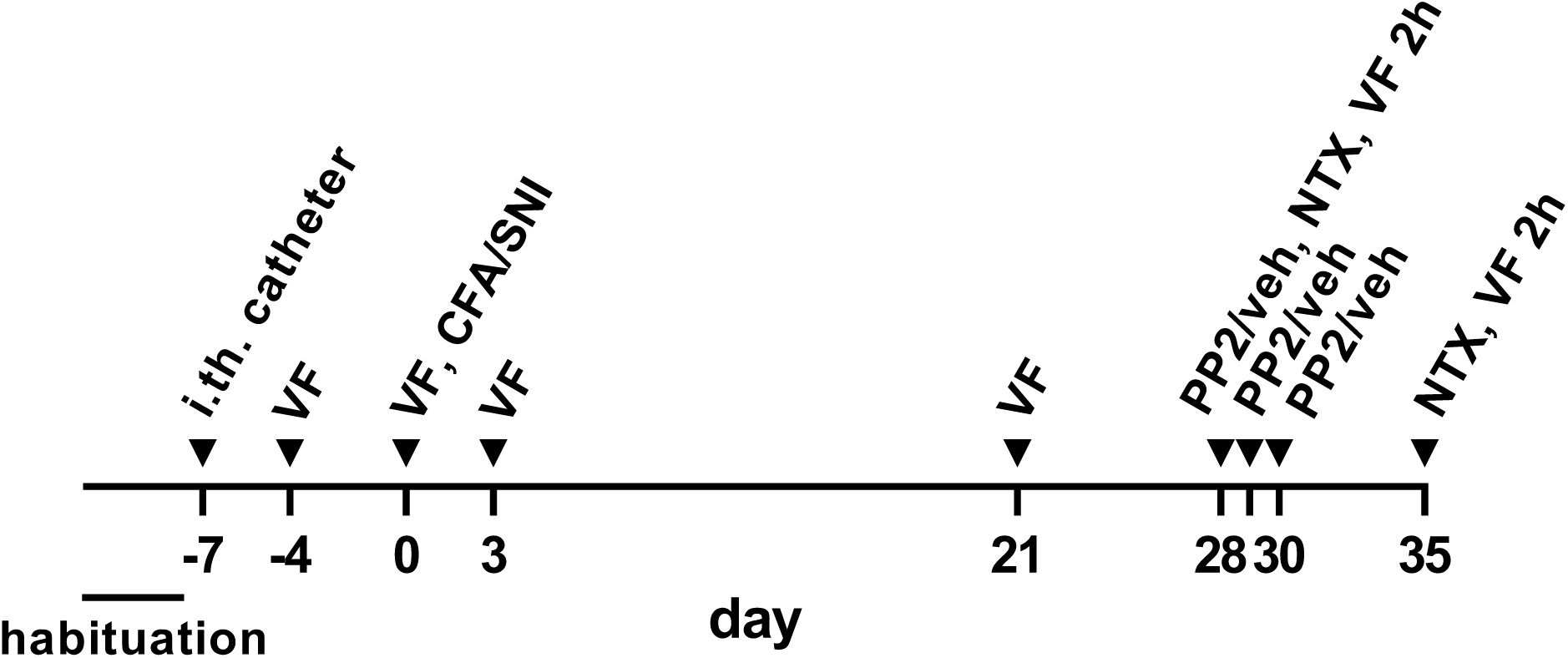
Timeline of the experiments: habituation - 30 min daily habituation to the von Frey cages for 3 days; i.th. catheter - intrathecal catheter implantation; VF - von Frey measures; VF 2h - von Frey measures for 2 h; CFA - 50 µl s.c. complete Freund’s adjuvant injected in the paw; SNI - spared nerve injury; PP2 - 0.3 nmol PP2 i.th. in 10 µl; veh - 10 µl vehicle i.th.; NTX - 2.6 nmol naltrexone i.th. in 10 µl. CFA, SNI, vehicle or PP2 were given to four different groups of rats.

### Sample size, randomization and blinding

The target sample size was *n* = 7-8 rats per group based on a power analysis, but 3 rats in the SNI-control group were excluded after day 28 because they had lost the i.th. catheter. Rats were randomly assigned to treatment and drug. For blinding, solutions of vehicle (for controls) and PP2 were prepared and randomly coded by JCM. WC injected the coded drugs i.th. and performed the von Frey measures. JCM analyzed the data and matched the codes with the drugs.

### Data analysis

Data were analyzed using Prism 8.3 (GraphPad Software, San Diego, CA) and expressed as mean ± standard error of the mean. Statistical significance was set at 0.05. Statistical analyses consisted of repeated measures (by both factors) two-way ANOVA followed by Holm-Sidak’s post-hoc tests.

## Results

### Latent sensitization induced by CFA

To determine if SFKs mediate the expression and maintenance of latent sensitization, we studied the short-term and long-term effects of the SFK inhibitor PP2 on the mechanical allodynia induced by NTX in rats with latent sensitization. One form of latent sensitization was induced by injecting 50 µl CFA in one hindpaw, a model of inflammatory chronic pain. Two groups of 8 and 7 rats were treated following the timeline shown in Fig. 1. In both groups, CFA produced mechanical allodynia in the ipsilateral but not in the contralateral hindpaw (Fig. 2A, D). Ipsilateral allodynia was strong on day 3 and decreased to near baseline values on days 21 and 28 (Fig. 2A, D). On day 28, rats were injected i.th. with vehicle (0.3% DMSO in saline) or PP2 (0.3 nmol) followed 15 min later by i.th. NTX (2.6 nmol). In the vehicle-injected rats, NTX induced mechanical allodynia in both the ipsilateral and the contralateral hindpaws, which peaked at 15 min and largely disappeared at 2 h (Fig. 2B). NTX had no effect in mice or rats injected with saline in the paw instead of CFA (Corder et al., 2013; Walwyn et al., 2016). NTX-induced bilateral hyperalgesia is a hallmark of latent sensitization (Campillo et al., 2011; Corder et al., 2013; Marvizon et al., 2015; Walwyn et al., 2016). In contrast, NTX did not induce allodynia after PP2 (Fig. 2E). Vehicle or PP2 were injected daily for two more days (Fig. 1). On day 35, rats received a second i.th. injection of NTX. In the control group, NTX again induced bilateral allodynia that peaked at 15 min and was gone after 2 h (Fig. 2C). In the rats injected with PP2, NTX-induced allodynia was absent on day 35 (Fig. 2F).

**Figure 2.**
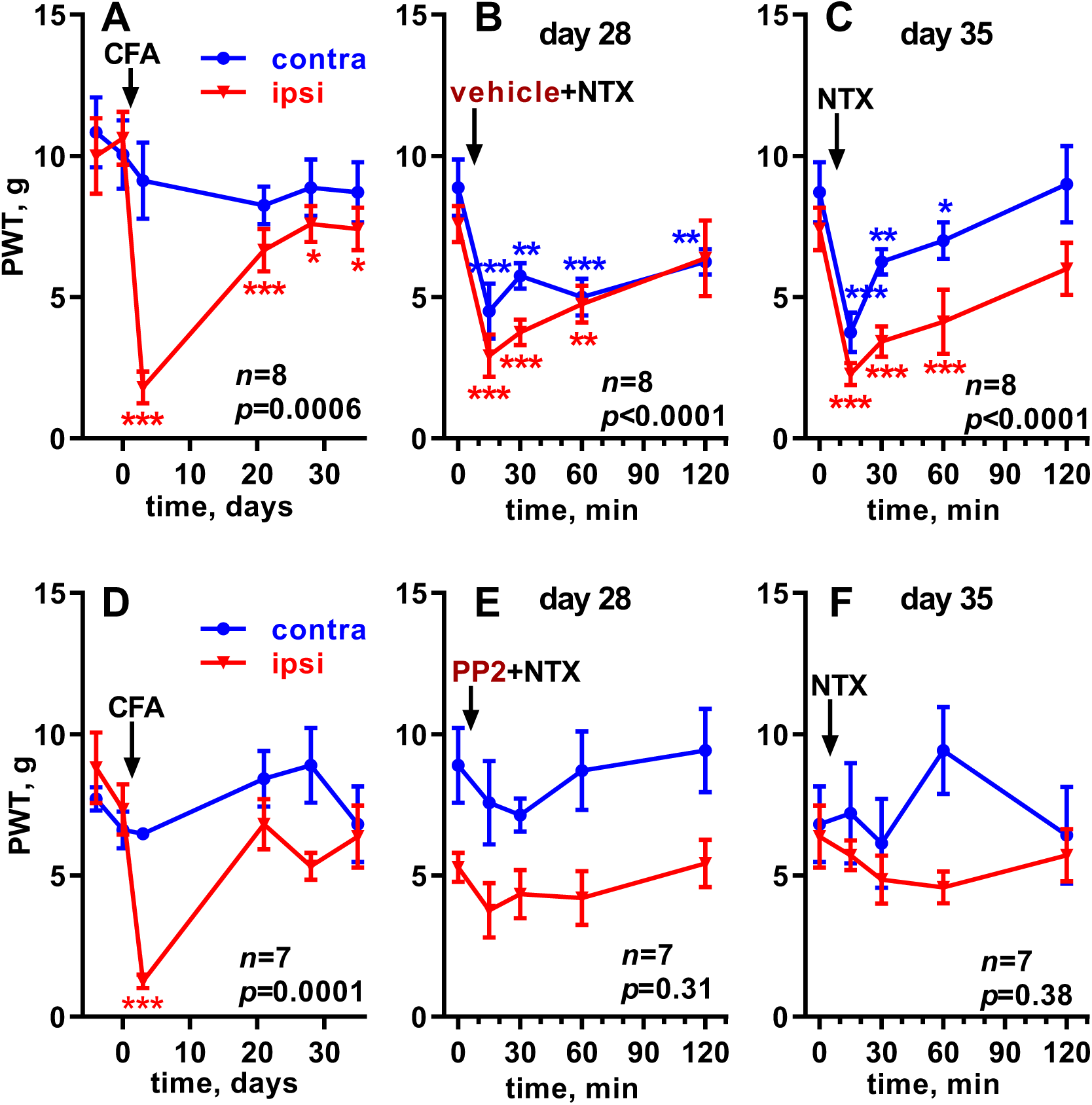
Effect of PP2 on NTX-induced allodynia in rats with CFA-induced latent sensitization. **A, D:** Rats received 50 µl CFA in one hindpaw. Paw withdrawal thresholds (PWT, von Frey filaments) were measured in the ipsilateral (‘ipsi’) and contralateral (‘contra’) hindpaws. **B, E:** On day 28, rats received i.th. injections of 10 µl vehicle (0.3% DMSO in saline, or PP2 (0.3 nmol in 0.3% DMSO, E), and 15 min later 10 µl NTX (2.6 nmol). **C, F:** Vehicle or PP2 were injected i.th. daily for two more days. On day 35, rats received i.th. injections of 10 µl NTX (2.6 nmol). Each panel was analyzed by repeated-measures (both variables) 2-way ANOVA; *p* values for ‘time’ (effect of CFA or NTX) are given in each panel, full results are given in Table 1. Holm-Sidak’s post-hoc tests comparing to time 0: * *p*<0.05, ** *p*<0.01, *** *p*<0.001.

Results in Fig. 2 were analyzed by 2-way ANOVA with repeated measures for the two variables: time and hindpaw side (Table 1). Note that the variable ‘time’ indicates whether there was a significant effect of CFA (rows A and D) or NTX. As expected, CFA produced a significant effect over time in both the control (row A) and the PP2 groups (row D), with significant differences between the paw sides. The effect of NTX over time was significant in control rats (rows B, C) but not in rats injected with PP2 (rows E, F). The interaction of the variables ‘time’ and ‘side’ (‘time x side’) was significant after the injection of CFA but not after the injections of NTX, indicating that CFA produced allodynia ipsilaterally while NTX produced allodynia bilaterally.

**Table 1.**
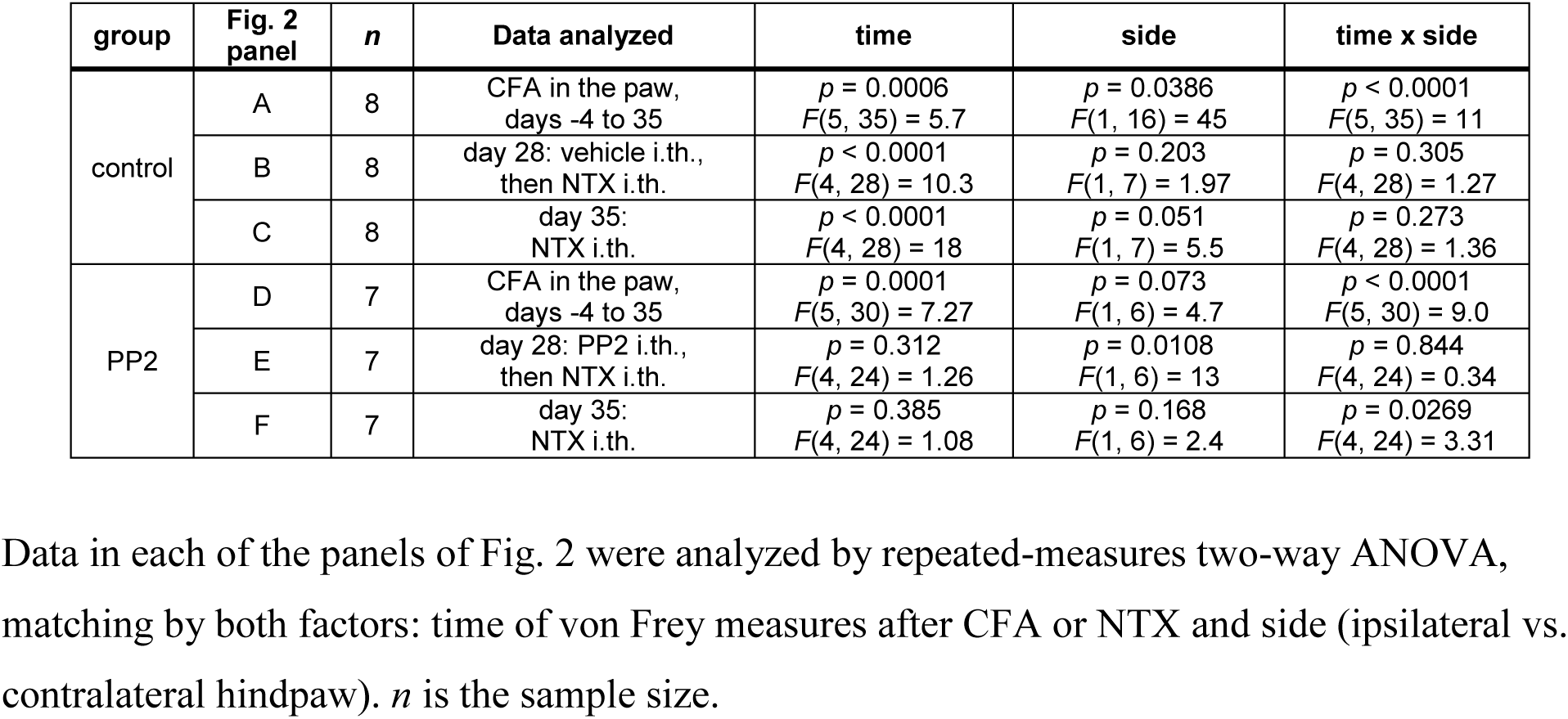
Repeated measures two-way ANOVA of CFA-induced LS data in Fig. 2

| group | Fig. 2 panel | *n* | Data analyzed | time | side | time x side |
| --- | --- | --- | --- | --- | --- | --- |
| control | A | 8 | CFA in the paw, days -4 to 35 | $p = 0.0006$<br>$F(5, 35) = 5.7$ | $p = 0.0386$<br>$F(1, 16) = 45$ | $p < 0.0001$<br>$F(5, 35) = 11$ |
| | B | 8 | day 28: vehicle i.th., then NTX i.th. | $p < 0.0001$<br>$F(4, 28) = 10.3$ | $p = 0.203$<br>$F(1, 7) = 1.97$ | $p = 0.305$<br>$F(4, 28) = 1.27$ |
| | C | 8 | day 35: NTX i.th. | $p < 0.0001$<br>$F(4, 28) = 18$ | $p = 0.051$<br>$F(1, 7) = 5.5$ | $p = 0.273$<br>$F(4, 28) = 1.36$ |
| PP2 | D | 7 | CFA in the paw, days -4 to 35 | $p = 0.0001$<br>$F(5, 30) = 7.27$ | $p = 0.073$<br>$F(1, 6) = 4.7$ | $p < 0.0001$<br>$F(5, 30) = 9.0$ |
| | E | 7 | day 28: PP2 i.th., then NTX i.th. | $p = 0.312$<br>$F(4, 24) = 1.26$ | $p = 0.0108$<br>$F(1, 6) = 13$ | $p = 0.844$<br>$F(4, 24) = 0.34$ |
| | F | 7 | day 35: NTX i.th. | $p = 0.385$<br>$F(4, 24) = 1.08$ | $p = 0.168$<br>$F(1, 6) = 2.4$ | $p = 0.0269$<br>$F(4, 24) = 3.31$ |

### Latent sensitization induced by SNI

We also studied the effect of PP2 on the latent sensitization induced by nerve injury. SNI was performed by cutting the common peroneal and sural branches of the sciatic nerve (CpxSx) (Solway et al., 2011) instead of the common peroneal and tibial nerves (Marvizon et al., 2019), because CpxSx produces allodynia that lasts only ∼28 days and not several months like other nerve injury models. This allowed us to test the effect of NTX after responses returned to baseline.

Two groups of 8 and 7 rats were treated following the timeline shown in Fig. 1. In both groups, CpxSx SNI produced mechanical allodynia in the ipsilateral but not in the contralateral hindpaw (Fig. 3A, D). Ipsilateral allodynia was strong on day 3 and decreased to near baseline values by day 28 (Fig. 3A, D). On day 28, the rats were injected i.th. with vehicle (0.3% DMSO in saline) or PP2 (0.3 nmol) and 15 min later with NTX (2.6 nmol). In the control group, NTX induced mechanical allodynia in both the ipsilateral and the contralateral hindpaws, which peaked at 30 min and, at 2 h, decreased ipsilaterally and overshot the baseline contralaterally (Fig. 3B). In contrast, NTX did not induce allodynia after PP2, with values that overshot the baseline at 2 h (Fig. 3E). Vehicle or PP2 were injected daily for two more days (Fig. 1). On day 35 rats received a second injection of NTX. In the control group, NTX again induced bilateral allodynia that peaked at 30 min and was gone after 2 h (Fig. 3C). In the rats injected with PP2, NTX-induced allodynia was absent on day 35 (Fig. 3F).

**Figure 3.**
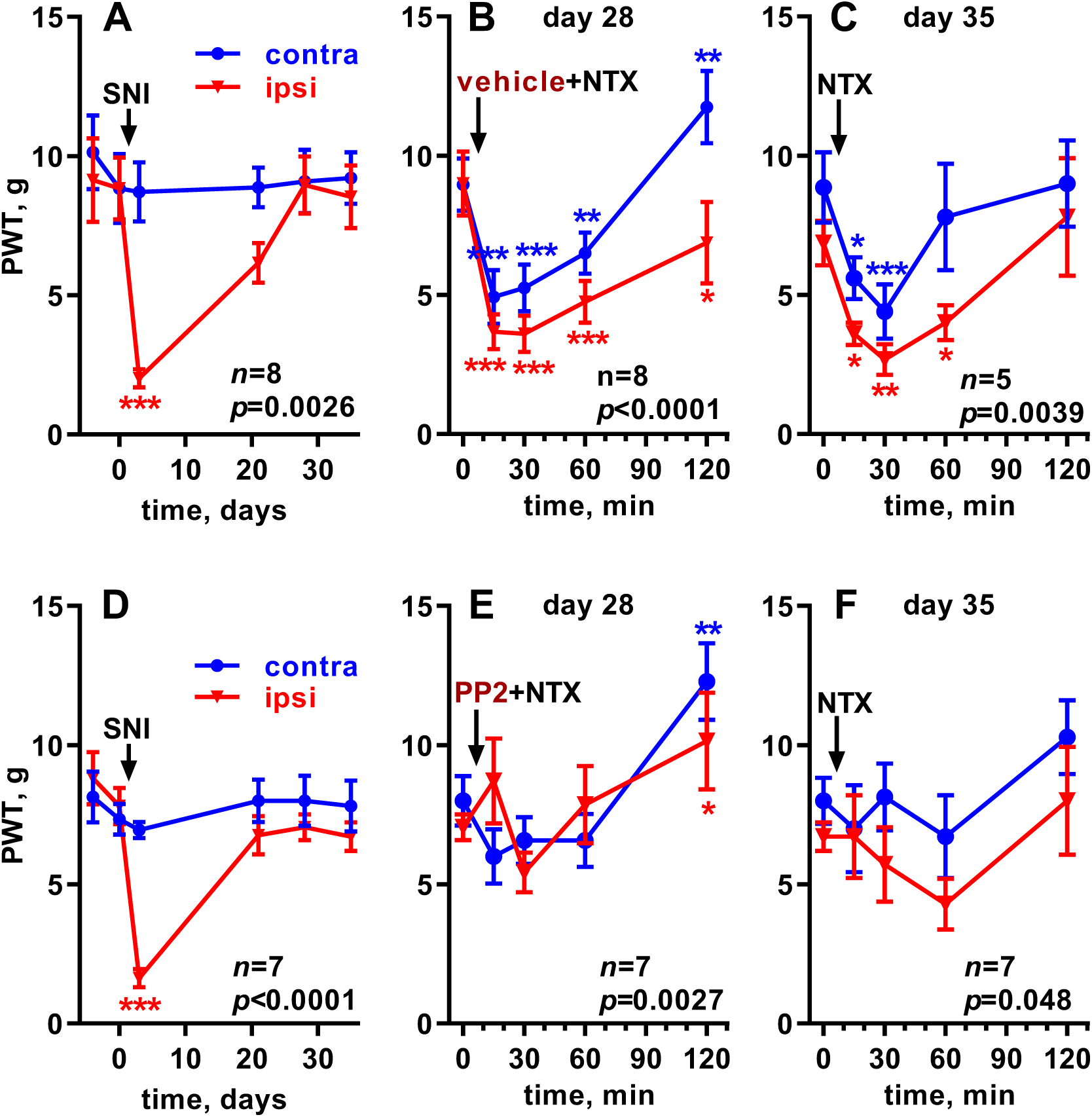
Effect of PP2 on NTX-induced allodynia in rats with SNI-induced latent sensitization. **A, D:** Rats received SNI in one hindpaw. Paw withdrawal thresholds (PWT, von Frey filaments) were measured in the ipsilateral (‘ipsi’) and contralateral (‘contra’) hindpaws. **B, E:** On day 28, rats received i.th. injections of 10 µl vehicle (0.3% DMSO in saline, B) or PP2 (0.3 nmol in 0.3% DMSO, E), and 15 min later 10 µl NTX (2.6 nmol). **C, F:** Vehicle or PP2 were injected i.th. daily for two more days. On day 35, rats received i.th. injections of 10 µl NTX (2.6 nmol). Each panel was analyzed by repeated-measures (both variables) 2-way ANOVA; *p* values for ‘time’ (effect of CFA or NTX) are given in each panel, full results are given in Table 2. Holm-Sidak’s post-hoc tests comparing to time 0: * *p*<0.05, ** *p*<0.01, *** *p*<0.001.

Table 2 shows the results of a 2-way ANOVA, repeated measures by both variables, of the data in Fig. 3. As expected, SNI produced a significant effect over time in both the control (row and the PP2 groups (row D). There were significant differences between sides and significant interaction ‘time x side’, showing that SNI produced ipsilateral allodynia. On days 28 and 35, NTX produced a significant effect over time in the control rats (rows B, C). In the rats injected with PP2, the effect of NTX was also significant on day 28 (row E), but this was due to the decrease in allodynia at the 120 min time point (Fig. 3E) and not to an increase in allodynia at 30 min. On day 35 in the PP2-injected rats, the effect of NTX was marginally significant (row F) probably due to a small increase in allodynia in the ipsilateral paw at 60 min and a small decrease in allodynia at 120 min in the contralateral paw (Fig. 3F). However, these effects were not significant in the Holm-Sidak’s post-hoc test. Overall, the allodynia produced by NTX in rats with SNI-induced latent sensitization was eliminated by PP2.

**Table 2.** Repeated measures two-way ANOVA of SNI-induced LS data in Fig. 3

| group | Fig. 3 panel | *n* | Data analyzed | time | side | time x side |
| --- | --- | --- | --- | --- | --- | --- |
| control | A | 8 | SNI, days -4 to 35 | $p = 0.0026$<br>$F(5, 35) = 4.58$ | $p = 0.0017$<br>$F(1, 7) = 24$ | $p < 0.0001$<br>$F(5, 29) = 8.3$ |
| | B | 8 | day 28: vehicle i.th., then NTX i.th. | $p < 0.0001$<br>$F(4, 28) = 16$ | $p = 0.0085$<br>$F(1, 7) = 13$ | $p = 0.0042$<br>$F(4, 28) = 4.8$ |
| | C | 5 | day 35: NTX i.th. | $p = 0.0039$<br>$F(4, 16) = 6.0$ | $p = 0.063$<br>$F(1, 4) = 27$ | $p = 0.403$<br>$F(4, 16) = 1.07$ |
| PP2 | D | 7 | SNI, days -4 to 35 | $p < 0.0001$<br>$F(5, 30) = 12$ | $p = 0.013$<br>$F(1, 6) = 12$ | $p = 0.0028$<br>$F(5, 30) = 4.7$ |
| | E | 7 | day 28: PP2 i.th., then NTX i.th. | $p = 0.0027$<br>$F(4, 24) = 5.5$ | $p = 0.901$<br>$F(1, 6) = 0.016$ | $p = 0.029$<br>$F(4, 24) = 3.2$ |
| | F | 7 | day 35: NTX i.th. | $p = 0.048$<br>$F(4, 24) = 2.8$ | $p = 0.0023$<br>$F(1, 6) = 26$ | $p = 0.723$<br>$F(4, 24) = 0.52$ |

## Discussion

This study shows that the SFK inhibitor PP2 eliminated the mechanical allodynia produced by NTX in rats with latent sensitization induced by CFA or SNI. A single i.th. injection of PP2 eliminated the effect of NTX given shortly afterward, whereas two more daily injections of PP2 eliminated the allodynia induced by NTX given five days after the last injection. This indicates that an SFK, likely Fyn, is involved in the maintenance of latent sensitization, so that when this SFK is inhibited latent sensitization disappears.

Previous studies indicate that the SFK Fyn forms part of a signaling pathway initiated by BDNF release from primary afferents (Lever et al., 2001) and microglia (Coull et al., 2005; Ulmann et al., 2008). BDNF binds to trkB receptors in primary afferents (Chen et al., 2014; Li et al., 2017) and dorsal horn neurons (Geng et al., 2010), which can activate Fyn through two pathways, one involving phosphoinositide 3-kinase and protein kinase B and the other mediated by Raf, MEK1/2 and ERK (Xu et al., 2006; Geng et al., 2010; Hildebrand et al., 2016; Marcos et al., 2017). Fyn then induces the activating phosphorylation of NMDA receptors (Guo et al., 2002; Abe et al., 2005; Chen et al., 2010), increasing substance P release (Chen et al., 2014), long-term potentiation of synapses in nociceptive pathways (Li et al., 2017) and neuropathic pain (Geng et al., 2010; Katano et al., 2011; Li et al., 2011). We have shown that BDNF induces the phosphorylation of NMDA receptors in primary afferents at Tyr^1472^ of their NR2B subunit, the site phosphorylated by Fyn (Li et al., 2017), and that this increases NMDA receptor currents and NMDA receptor-induced substance P release (Chen et al., 2014). Substance P activates neurokinin 1 receptors in lamina I neurons, which project to the parabrachial nucleus (Todd et al., 2000; Todd et al., 2002) and are required for latent sensitization induced by surgical incision and opioids (Rivat et al., 2009; Arora et al., 2016).

In contrast with our results, Araldi et al. (2017; 2018) found that the SFK inhibitor SU-6656 did not inhibit CFA-induced latent sensitization at the relatively high dose of 27 nmol i.th. However, they found that SU-6656 did inhibit latent sensitization when combined with the MEK1/2 inhibitor U0126 (24 nmol i.th.). The effect for PP2 described here was obtained at a dose 100 times lower than the dose of SU-6656 used by Araldi et al., even though PP2 and SU-6656 have similar potencies to inhibit SFKs (Lawrence and Niu, 1998; Bain et al., 2003). However, we cannot rule out that PP2 and SU-6656 have different effects on Fyn. Thus, if Fyn gets activated by BDNF through the Raf - MEK1/2 - ERK pathway, complete inhibition of Fyn by PP2 may make it unnecessary the additional inhibition of MEK1/2 by U0126.

Araldi et al. (2017; 2018) also argued that latent sensitization is the same phenomenon as hyperalgesic priming type 2, which was also blocked by co-administration of the SFK inhibitor SU-6656 and the MEK1/2 inhibitor U0126. Hyperalgesic priming type 2 is induced by repeated intradermal injection of a µ-opioid receptor agonist (Araldi et al., 2015, 2017). Although a form of latent sensitization is induced by overdoses of opioids like fentanyl and remifentanil, its hyperalgesia is additive with the hyperalgesia of latent sensitization induced by paw incision (Campillo et al., 2011) or carrageenan (Rivat et al., 2007; Laboureyras et al., 2014). This suggests that opioid-induced latent sensitization is different from the latent sensitization induced by inflammation, incision or nerve injury. Therefore, there is no reason at present to believe that latent sensitization is the same phenomenon as hyperalgesic priming type 2. As for hyperalgesic priming type 1 (Parada et al., 2005; Joseph and Levine, 2010; Ferrari et al., 2013), it has properties that are quite different from latent sensitization (Marvizon et al., 2015).

An odd characteristic of latent sensitization is that, while the initial mechanical allodynia produced by CFA or SNI occurs only in the ipsilateral paw, the allodynia induced by NTX and other opioid antagonists occurs bilaterally (Campillo et al., 2011; Corder et al., 2013; Marvizon et al., 2015; Walwyn et al., 2016). Bilateral mechanical allodynia was also induced by a spinal block and by i.c.v. CRF in rats with latent sensitization (Chen et al., 2018), indicating that the sensitization spreads to the contralateral side of the spinal cord and is suppressed bilaterally by ongoing descending pain inhibition. Contralateral or “mirror pain” is a well-established phenomenon that has been attributed to astrocyte activation (Sluka et al., 2001; Gao et al., 2010). Here we show that PP2 is able to eliminate both the ipsilateral and the contralateral sensitization.

One caveat of our approach is that PP2 is not entirely selective for SFKs (Lawrence and Niu, 1998; Bain et al., 2003). However, PP2 has been used in most studies targeting Fyn. Future studies are needed to determine if the BDNF signaling pathway leading to Fyn activation proposed here is the one that maintains latent sensitization. Our preliminary results show that the BDNF scavenger trkB-Fc (Li et al., 2017) also eliminates NTX-induced allodynia in rats with SNI-induced latent sensitization.

There is evidence that BDNF mediates neuropathic pain by downregulating the KCC2 cotransporter, which increases intracellular Cl^-^ and thus cancels the inhibition by GABA_A_ and glycine receptors of excitatory interneurons (Coull et al., 2005; Gagnon et al., 2013). Inflammatory chronic pain appears to also be mediated by BDNF (Lin et al., 2011; Lalisse et al., 2018), although it is still unclear if different mechanisms mediate inflammatory and neuropathic pain (Peirs et al., 2015; Peirs and Seal, 2016). Our results show that an SFK is involved in the maintenance of latent sensitization induced by both inflammation and nerve injury. It will be important to elucidate if BDNF is involved in these two forms of latent sensitization and whether they have the same maintenance mechanisms.

## Acknowledgements

This study was done under the umbrella of the following UCLA institutes: Brain Research Institute, Center for the Study of Opioid Receptors and Drugs of Abuse, CURE: Digestive Diseases Research Center, and the Oppenheimer Family Center for Neurobiology of Stress and Resilience.

## Authors contributions

W. Chen performed the surgeries, obtained behavioral measures and contributed to the design of the study. J.C. Marvizón designed the study, analyzed the data, made the figures and wrote the manuscript.

## Data accessibility

Data will be archived in Figshare.

## Abbreviations

ANOVA: analysis of variance
BDNF: brain-derived neurotrophic factor
CFA: complete Freund’s adjuvant
ERK1: extracellular signal-regulated kinase 1
Fig: Figure
i.th: intrathecal
MEK1/2: mitogen-activated protein kinase kinases 1 and 2 NTX naltrexone
NMDA: N-methyl-D-aspartate
PP2: 3-(4-chlorophenyl) 1-(1,1-dimethylethyl)-1H-pyrazolo[3,4-d]pyrimidin-4-amine
PWT: paw withdrawal threshold

